# UBL3 UBL domain exhibits distinct helix-centered dynamic control among ubiquitin-like proteins

**DOI:** 10.64898/2026.04.06.716645

**Authors:** Kazuki Matsuda, Yuko Moriya, Lili Xu, Ryo Ohmagari, Shuhei Aramaki, Chi Zhang, Atsushi Baba, Shoshiro Hirayama, Tomoaki Kahyo, Mitsutoshi Setou

## Abstract

Ubiquitin-like protein 3 (UBL3) is a post-translational modifier that sorts proteins into small extracellular vesicles and regulates the trafficking of disease-associated proteins such as α-synuclein. The structural and dynamic features of the UBL domain that underlie these functions, however, remain poorly understood. Here we performed in silico structural dynamics analysis of the UBL3 UBL domain using an NMR structure ensemble combined with anisotropic network modeling (ANM) and perturbation response scanning (PRS). Principal component analysis and residue-wise fluctuation analysis consistently revealed high flexibility in the C-terminal region of UBL3. Comparative ANM analysis across 20 ubiquitin-like proteins (UBLs) further showed that C-terminal flexibility is a conserved yet variable property within the UBL family. PRS analysis demonstrated that residues forming the central α-helix of the β-grasp fold exert greater dynamic control over collective motions than β-sheet residues. Notably, UBL3 displayed the highest helix/sheet PRS effectiveness ratio among all UBLs analyzed, highlighting the prominent dynamic contribution of helix residues in this domain. Together, these results provide a structural basis for understanding UBL3-dependent protein interactions and disease-related mechanisms, and suggest that helix-centered dynamic control in the UBL domain may represent a potential target for modulating UBL3 function.

## INTRODUCTION

UBL3 (Ubiquitin-like 3) was first identified in Drosophila in 1999 [1]. This protein is highly conserved across eukaryotes and anchors to the plasma membrane via prenylation at a C-terminal CAAX motif [2]. Functional analyses revealed that UBL3, unlike canonical ubiquitin, acts as a novel post-translational modifier (PTM) that modifies target proteins through a terminal cysteine residue; it also localizes to extracellular vesicles [3][4]. This UBL3 modification system is thought to serve as a major protein-sorting mechanism, on par with the ubiquitin–proteasome and autophagy systems, directing proteins into small extracellular vesicles (sEVs) [5]. Intracellularly, UBL3 undergoes dynamic transport from the cytoplasm shortly after synthesis to the plasma membrane and multivesicular bodies (MVBs), ultimately accumulating in MVBs [6]. In certain cell types, UBL3 also traffics to primary cilia via PDE6D [7]. In the immune system, UBL3 functions as an essential regulator of MARCH1-dependent trafficking of MHC II and CD86 in dendritic cells [8].

Regarding disease associations, UBL3 acts as a tumor suppressor in non-small cell lung cancer [9]. In pancreatic cancer, it has been identified as a biomarker that predicts patient survival [10]. Its utility as a biomarker for cancer progression has also been identified in gastric cancer [11]. In head and neck squamous cell carcinoma, low UBL3 expression contributes to poor prognosis [12]. More recently, statins were reported to enhance cancer immunotherapy efficacy by inhibiting UBL3 modification [13]. UBL3 also regulates target cell survival during articular cartilage repair [14]. In the context of neurodegeneration, UBL3 directly interacts with α-synuclein, which is implicated in Parkinson’s disease [15]. This interaction is positively regulated by the antioxidant enzyme MGST3 [16][17]. In Huntington’s disease, UBL3 interacts with mutant huntingtin fragments and modifies their intracellular sorting [18]. UBL3 has also been shown to interact with RNA-binding proteins such as FUS [19].

Thus, UBL3 is known to interact with an exceptionally diverse set of target proteins. How UBL3 recognizes and engages such a broad range of interacting partners through its structural dynamics, however, remains poorly understood, and the structural dynamics of the UBL domain that forms the functional core of UBL3 have not been systematically analyzed.

Structurally, UBL3 belongs to the ubiquitin-like protein (UBL) superfamily. UBLs share the β-grasp fold exemplified by ubiquitin as their basic scaffold, and this evolutionarily conserved structural motif underpins diverse cellular functions [20]. The β-grasp fold—comprising a five-stranded β-sheet and a central α-helix—has been reused as a stable core platform for post-translational modifiers and interaction modules [21]. Within the UBL family, however, notable diversity exists in surface charge distribution, loop regions, α-helix arrangement, and the sequence and flexibility of terminal regions; these differences are recognized as key determinants of binding partner specificity [22]. For example, SUMO-1 and SUMO-2/3 display distinct interaction properties in non-covalent binding to UBL domains via SUMO-interacting motifs (SIMs) [23]. In the ATG8 family (LC3s/GABARAPs), each isoform shows functional diversity in cells through differences in binding selectivity for LIR/AIM motifs present in autophagy-related target proteins, despite sharing a conserved β-grasp fold [24]. Bacterial ubiquitin-like proteins, too, have been reported to display multi-domain architectures and oligomerization based on the β-grasp scaffold, revealing diverse modes of structural repurposing [25]. Taken together, UBLs are understood to have acquired diverse biological functions through local structural plasticity while retaining a common three-dimensional scaffold. For UBL3 specifically, unique functions including CAAX-dependent prenylation, membrane anchoring, and atypical cysteine-dependent modification have been established, yet the structural features and dynamic properties of the underlying UBL domain have not been systematically analyzed [2]. Structural analysis focused on the UBL3 UBL domain is therefore important for linking the structural principles common to the UBL family with the functional properties unique to UBL3.

In protein kinases and PDZ domains, for example, central α-helices act as mechanical hubs that control long-range information transfer and global collective motions, as demonstrated by Perturbation-Response Analysis [26][27].

Chemical manipulation and modulation of UBLs has been shown to contribute to drug discovery strategies through chemical biology approaches [28]. Identifying structural elements that determine interaction selectivity or serve as regulatory points via the UBL domain is therefore essential for understanding, at the molecular level, interactions with disease-relevant proteins that could inform therapeutic development. In the present study, we focused on the UBL domain of UBL3 and aimed to characterize its structural features through in silico analysis using an NMR structure ensemble and structural prediction models, with particular attention to structural fluctuations and residue dynamics related to secondary structure.

## MATERIALS AND METHODS

### Structural Diversity Analysis

NMR-determined protein structure data were downloaded from the RCSB Protein Data Bank (https://www.rcsb.org/) and used for analysis; the datasets are summarized in Table 1. To evaluate structural diversity, we used EnsembleFlex [29]. Backbone atoms (N, Cα, C, O) were extracted from each PDB model, and structural alignment was performed using the pdbaln function. Structures were then superimposed by least-squares fitting of Cα atoms using rotation and translation operations, placing all models in a common coordinate frame. All subsequent analyses in this study used the superimposed backbone structures.

**Table 1.** Summary of NMR structures used in this study.

| Protein Name | PDB ID§ | UBL Type | Number of Models | UniProt ID (Total Residues) | PDB Range† | UniProt Range† | PDB UBL‡ | UniProt UBL‡ |
| --- | --- | --- | --- | --- | --- | --- | --- | --- |
| UBL3 | 2GOW | Standalone | 20 | O95164 (117) | 2–117 | 2–117 | 10–88 | 10–88 |
| UBIQUITIN (Ub) | 1D3Z | Standalone | 10 | P0CG48 (685) | 1–76 | 609–684 | 1–76 | 609–684 |
| SUMO1 | 2N1V | Standalone | 20 | P63165 (97) | 1–97 | 1–97 | 20–97 | 20–97 |
| SUMO2 | 2N1W | Standalone | 20 | P61956 (95) | 1–93 | 1–93 | 16–95 | 16–95 |
| SUMO3 | 1U4A | Standalone | 20 | P55854 (103) | 14–92 | 14–92 | 15–92 | 15–92 |
| NEDD8 | 2KO3 | Standalone | 20 | Q15843 (76) | 1–76 | 1–76 | 1–76 | 1–76 |
| LC3B | 1V49 | Standalone | 1 | Q9GZQ8 (125) | 1–120 | 1–120 | 1–120 | 1–120 |
| UFM1 | 1WXS | Standalone | 10 | P61960 (85) | –4–83 | 1–85 | 1–83 | 1–83 |
| HUB1 | 1P0R | Standalone | 1 | Q9BZL1 (73) | 1–73 | 1–73 | 1–73 | 1–73 |
| ISG15 | 2HJ8 | Tandem | 20 | P05161 (165) | 86–160 | 79–157 | 86–160 | 79–157 |
| FAT10 | 6GF2 | Tandem | 10 | O15205 (165) | 1–85 | 85–165 | 10–83 | 6–81 |
| ZFAND1 | 8HW9 | Domain | 20 | Q8TCF1 (268) | 130–268 | 133–268 | 160–260 | 160–260 |
| DC-UBP | 1TTN | Domain | 15 | Q8WUN7 (234) | 21–100 | 129–234 | 24–99 | 152–227 |
| SF3A1 | 7P08 | Domain | 20 | Q15459 (793) | 701–793 | 704–793 | 707–793 | 707–793 |
| NFATC2IP | 2JXX | Domain | 20 | Q8NCF5 (419) | 20–97 | 342–419 | 26–97 | 348–419 |
| Ubiquilin-1 | 2KLC | Domain | 20 | Q9UMX0 (589) | 23–101 | 34–112 | 26–100 | 37–111 |
| Ubiquilin-2 | 1J8C | Domain | 20 | Q9UHD9 (624) | 1–103 | 1–117 | 33–107 | 33–107 |
| DDI2 | 2N7D | Domain | 40 | Q5TDH0 (399) | 80–176 | 1–76 | 101–176 | 1–81 |
| PRKN | 1IYF | Domain | 10 | O60260 (465) | 1–76 | 1–76 | 1–76 | 1–76 |
| HHR23A | 1P98 | Domain | 11 | P54725 (363) | 1–78 | 1–78 | 1–78 | 1–81 |

§ PDB entries 1U4A and 6GF2 correspond to mutant sequences that differ from the UniProt reference.
† Residue ranges correspond between PDB and UniProt numbering and were used for ANM RMSF analysis, except for UFM1, in which the PDB coordinates contain non-UniProt residues and do not exactly match the UniProt-annotated region.
‡ UniProt-defined UBL regions were mapped to PDB residue numbering and used for PRS effectiveness analysis; when PDB coordinates did not fully cover the UniProt-annotated region, only the available residues were used.

To assess pairwise structural similarity, we computed an RMSD (Root Mean Square Deviation) matrix based on superimposed Cα coordinates. The resulting RMSD matrix was visualized as a heatmap to capture distance relationships within the structural ensemble.

Principal component analysis (PCA) was performed on the superimposed Cα coordinates to extract the principal axes representing major structural variation, and principal component scores (PC1, PC2, PC3) were calculated for each model.

To quantify local structural fluctuations, we computed per-residue RMSF (Root Mean Square Fluctuation) for Cα atoms using the superimposed backbone ensemble. RMSF was calculated as the magnitude of positional deviation from the mean coordinate across the ensemble for each residue’s Cα atom. Visualization was performed using ChimeraX (version 1.10.1).

### Anisotropic Network Model (ANM)

To evaluate the dynamic properties of protein structures, we built anisotropic network models (ANMs) using the ProDy library (version 2.6.1) [30]. In the ANM, each PDB structure was represented as a network in which Cα atoms serve as nodes and pairs of Cα atoms within a cutoff distance are connected by isotropic springs; the corresponding Hessian matrix was constructed. Per-residue mean-square fluctuations were computed from the 20 lowest-frequency normal modes—selected to capture cooperative motions across the entire structure (independent basic motion patterns available to the protein)—and defined as the ANM-based RMSF (ANM RMSF).

To evaluate the relative bias of fluctuations at the C-terminus, we defined the index RCₖ using the ANM RMSF fᵢ as shown in the equation below. RCₖ is the ratio of the arithmetic mean ANM RMSF of the last k residues to the arithmetic mean ANM RMSF of all residues. Here, fᵢ is the ANM RMSF value for residue i, N is the total number of residues analyzed, and Cₖ denotes the set of the first k residues counted from the C-terminus.

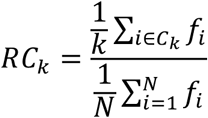

### Perturbation Response Scanning (PRS) Analysis

Perturbation Response Scanning (PRS) is a method that evaluates the dynamic network properties of proteins based on the propagation of responses when a perturbation is applied to individual residues [31]. Using the eigenvalues (mode stiffness) and eigenvectors (displacement directions) obtained from the ANM, we computed PRS effectiveness using the calcPerturbResponse function implemented in ProDy [32]:

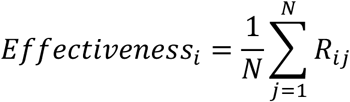

PRS effectiveness is defined as the mean response across all residues j when residue i is perturbed. Effectivenessᵢ is the PRS effectiveness of residue i. N is the number of residues analyzed, and Rᵢⱼ is the magnitude of the response of residue j to a perturbation at residue i, as computed by calcPerturbResponse.

Pocket geometry was characterized using vertex coordinates obtained from fpocket (version 4.0.2) as the pocket point cloud [33]. To compute solvent-accessible surface area (ASA), we used FreeSASA (version 2.2.1) to calculate per-atom ASA for all heavy atoms, then summed atomic ASA values within each residue to obtain per-residue ASA [34]. Pocket-contacting residues were defined as those for which the shortest distance between any atom in the residue and the pocket point cloud fell below a preset threshold (default: 5.0 Å) and whose residue ASA was greater than zero.

To evaluate PRS effectiveness by secondary structure, we ran DSSP (version 4.5.7) [35] and classified each residue in the UBL domain into three groups: Helix, Sheet, or Other. Between-group differences were assessed using Cliff’s δ as the effect size; correlations were assessed using Spearman’s rank correlation.

For comparison across UBLs, residues in the UBL domain were classified into Helix and Sheet groups. The helix/sheet PRS effectiveness ratio was then computed as follows:

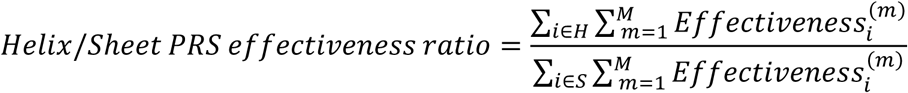

Here, H and S denote the sets of Helix and Sheet residues within the UBL domain, respectively. M and m represent the number of structural models (number of NMR models or Boltz2-generated candidates) and their index. Effectiveness_i_^(m)^ denotes the PRS effectiveness of residue i in structural model m.

Between-group differences were assessed using Cliff’s δ as the effect size. Uncertainty in Cliff’s δ was evaluated by computing 95% confidence intervals via bootstrap resampling (10,000 resamples). Correlations were assessed using Spearman’s rank correlation coefficient, with corresponding p-values calculated.

### Structure Prediction of UBL Domains Using Boltz2

Structure prediction was performed for the sequence regions annotated as UBL domains in UniProt. For standalone UBLs lacking a UniProt UBL domain annotation, the full-length sequence was used as the UBL domain.

Predictions were carried out using Boltz (version 2.2.1; Boltz2) [36][37]. For homology search, UniRef30 (release 2302) [38] and the ColabFold default environment database (sequences derived from clustering of the Big Fantastic Database and MGnify) were used [39], with MMseqs2 as the search algorithm [40]. The sensitivity parameter (-s) in MMseqs2 controls the trade-off between speed and sensitivity; we set it to 8.5 to enable high-sensitivity homology search. The maximum number of hits was set to 10,000 sequences from each database. The pairing strategy used server default settings. A multiple sequence alignment (MSA) was generated from the retrieved homologous sequences using the ColabFold API server (https://api.colabfold.com). The MSA was then provided as evolutionary input to Boltz2, and 10 candidate structures per sequence were generated using the default network weights (recycling steps = 20, seed = 42).

pLDDT is a per-residue confidence score introduced by AlphaFold [41]. Because this study involved single-chain structure prediction with Boltz2, we relied primarily on pLDDT for assessing prediction reliability [42]. All predicted structures exhibited sufficiently high pLDDT values and were therefore included in the analyses.

## RESULTS

### Fluctuation Analysis of the C-terminal Region of UBL3

The NMR protein structure dataset for UBL3 contains 20 structural models. Before characterizing structural features, we first analyzed positional differences among these models. Pairwise structural similarity was estimated using RMSD (Root Mean Square Deviation); no clear clusters were apparent from the heatmap (Figure 1A). The boundaries between models were not sharp: PCA also revealed a continuous data distribution (Figure 1B). Superposition of three representative models corresponding to the minimum, median, and maximum PC1 scores (models 12, 18, and 2, respectively) showed pronounced positional differences at the C-terminal region (Figure 1C). Per-residue contributions to PC1 were highest at the C-terminal region of UBL3 (Figure 1D). These results suggested that the C-terminal region of UBL3 adopts a flexible structure. To quantitatively verify this flexibility, we computed per-residue RMSF across structural models; the C-terminal region of UBL3 showed the highest values (Figure 1E). The ANM—constructed per model with Cα atoms as nodes and inter-node interactions approximated by springs, to evaluate collective motions and anisotropic fluctuations—also yielded high values at the C-terminal region (Figures 1F and 1G). Together, these in silico analyses consistently demonstrate that the C-terminal region of UBL3 is a region of high structural fluctuation.

**Figure 1.**
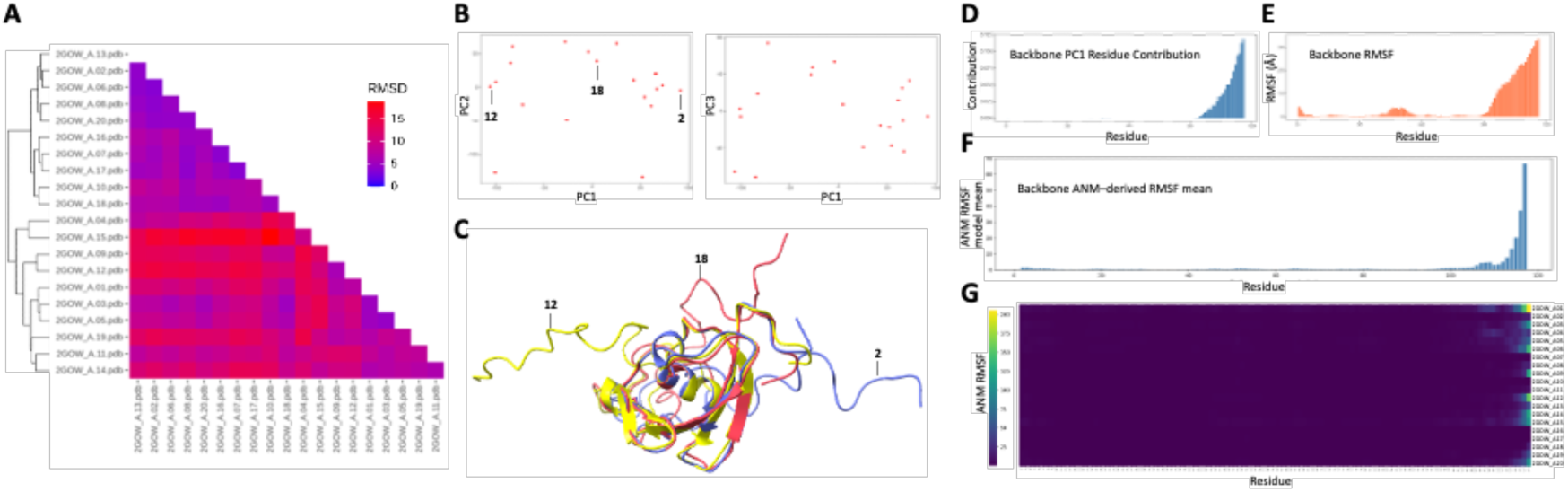
C-terminal region of UBL3 exhibits enhanced flexibility in the NMR ensemble. (A) Heatmap of pairwise Cα RMSD values calculated among the 20 NMR models of UBL3 after backbone superposition. (B) Scatter plots of principal component analysis (PCA) based on superimposed Cα coordinates, displaying continuous distributions along PC1–PC3. (C) Superposition of three representative models corresponding to the minimum, median, and maximum PC1 scores (models 12, 18, and 2, respectively), highlighting pronounced positional differences at the C-terminal region. (D) Residue-wise contributions to PC1. (E) Backbone RMSF calculated from the NMR ensemble. (F) Mean ANM-derived RMSF per residue calculated from the lowest 20 normal modes. (G) Heatmap of ANM-derived RMSF values across individual models, confirming consistently high flexibility at the C-terminal region.

### Fluctuation Analysis of C-terminal Regions Across UBLs

To determine whether high C-terminal fluctuation is specific to UBL3 or a broader property of the UBL family, we selected 20 UBLs—spanning standalone and domain types—for which NMR protein structure data were available. For each UBL, we computed the mean ANM RMSF of the last 1 to 10 C-terminal residues and derived the Relative C-terminal k-residue (RCk; k = 1–10) value as the ratio to the mean ANM RMSF across the entire UBL domain. An RCk value greater than 1 indicates that the last k residues fluctuate more than the protein-wide average. Within the C-terminal 1–10 residue window, 11 proteins including PRKN consistently showed high RCk values, while 5 proteins including NFATC2IP consistently showed low values (Figure 2). Among domain-type UBLs, PRKN and SF3A1 had relatively high RCk values; among standalone types, Ub, UBL3, and NEDD8 ranked highest compared with other UBLs. These results demonstrate that C-terminal fluctuation in the UBL domain, as assessed by ANM, is not unique to UBL3 but rather a variable structural property distributed across the UBL family.

**Figure 2.**
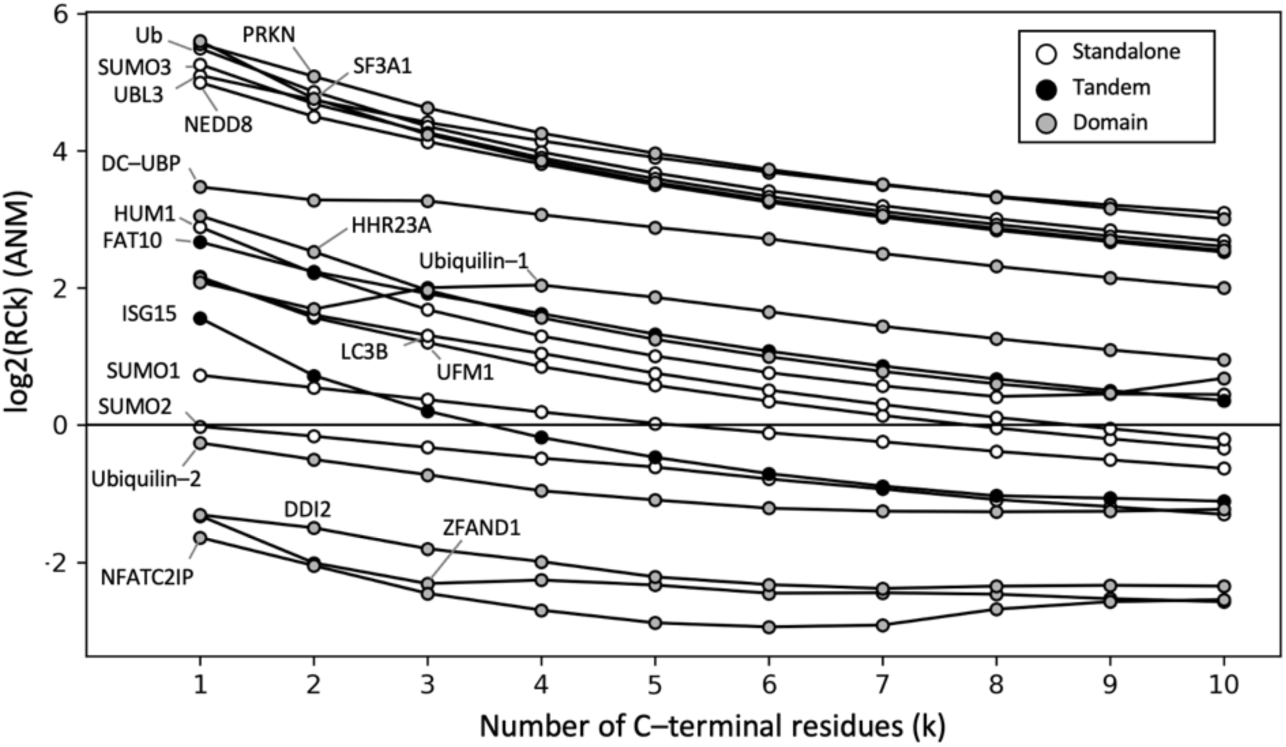
Comparative analysis of C-terminal fluctuations across ubiquitin-like proteins. Relative C-terminal k-residue (RCk; k = 1–10) values derived from ANM-based RMSF profiles and shown as log2-transformed values. RCk represents the ratio of the mean ANM RMSF of the last k residues to the mean ANM RMSF of all residues in the corresponding structure. Proteins are grouped as standalone, tandem, or domain-type ubiquitin-like proteins (UBLs).

### Secondary-Structure-Based Residue Perturbation Analysis in UBL3

We next computed PRS effectiveness for each residue in each UBL3 structural model, which quantifies how strongly each residue contributes to the large-amplitude, low-dimensional collective motions of the protein. Some UBL3 models showed very high PRS effectiveness at residues 32 to 36 (Figures 3A and 3B). Some of these residues contributed to pocket formation; when all residues were classified as pocket-contacting or non-pocket, pocket residues tended to show slightly higher PRS effectiveness (Cliff’s delta = 0.11) (Figure 3C). A moderate negative correlation was observed between pocket solvent-accessible surface area and PRS effectiveness (Spearman ρ = −0.42) (Figure 3D). Comparing PRS effectiveness among the Helix, Sheet, and Other groups, both Helix and Sheet residues showed higher values than Other residues. Effect size analysis revealed large effects for Helix vs. Other (Cliff’s δ = 0.414) and Sheet vs. Other (δ = 0.380), with a small difference also observed for Helix vs. Sheet (δ = 0.148) (Figure 3E). In particular, Ala32, Ala36, and Ser33—which ranked among the highest in mean PRS effectiveness across models—form the helix structure of the β-grasp fold, demonstrating a concentration of high-PRS-effectiveness residues in the helix of the UBL3 UBL domain (Figure 3F).

**Figure 3.**
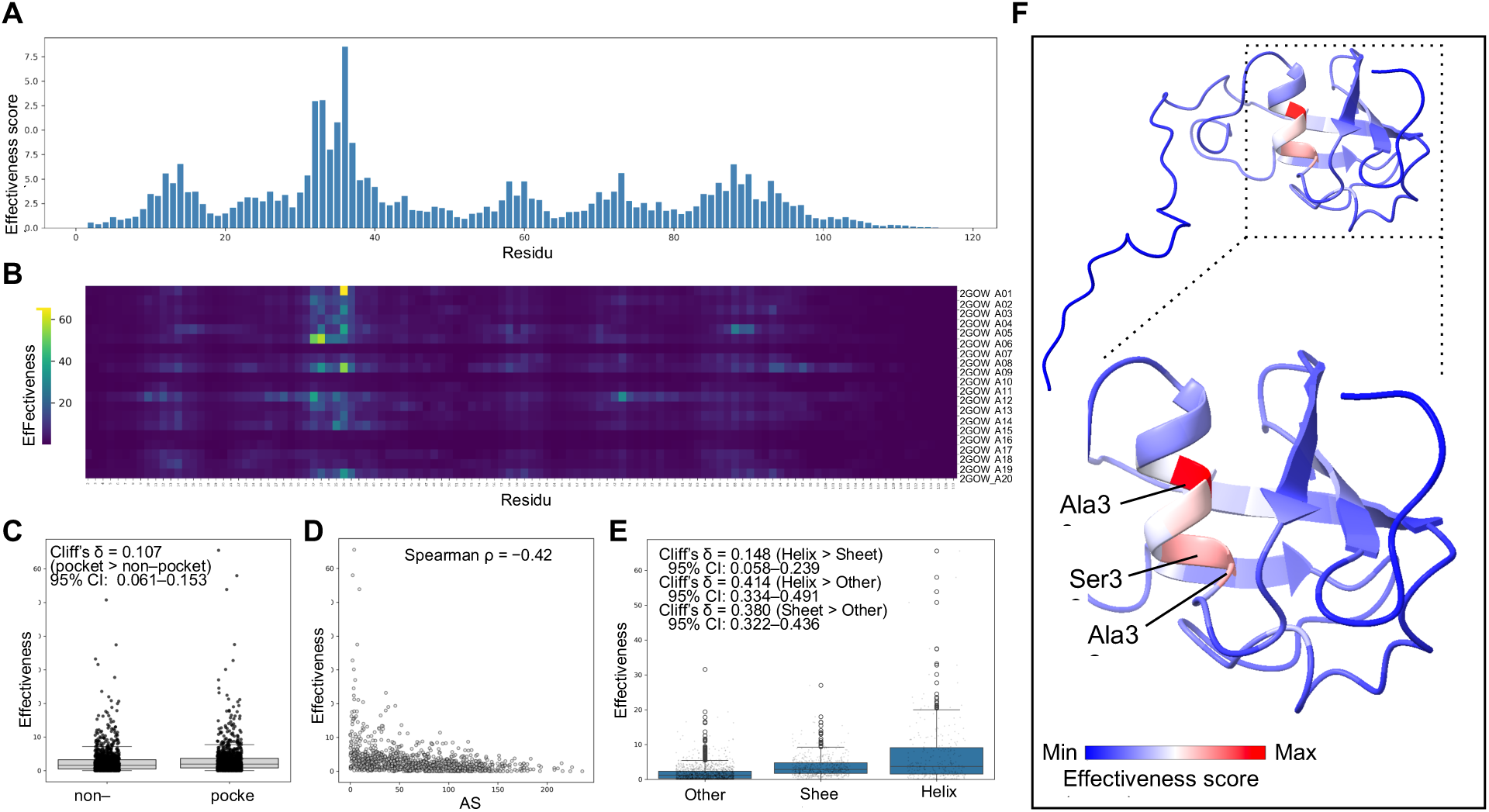
Perturbation response scanning reveals helix residues as major dynamic controllers in UBL3. (A) Residue-wise mean PRS effectiveness averaged across UBL3 NMR models. (B) Heatmap of PRS effectiveness for individual models. (C) Comparison of PRS effectiveness between pocket-contacting and non-pocket residues. (D) Scatter plot showing the relationship between residue solvent-accessible surface area (ASA) and PRS effectiveness. (E) Comparison of PRS effectiveness among residues classified as Helix, Sheet, or Other based on DSSP assignment, with effect sizes indicated. (F) Mapping of representative high-PRS-effectiveness residues (Ala32, Ser33, and Ala36) onto the UBL3 structure of model 1. Cliff’s δ values are shown with 95% bootstrap confidence intervals (10,000 resamples).

### Secondary-Structure-Based Residue Perturbation Analysis Across UBLs

To determine whether the concentration of high-PRS-effectiveness residues in helix structures is a feature shared among other UBLs, we compared the helix/sheet PRS effectiveness ratio across 20 UBL proteins. Based on NMR-derived structural data, UBL3 displayed the highest helix/sheet PRS effectiveness ratio among the 20 UBLs analyzed (Figure 4). Note that ratios below 1 are common because UBL domains generally contain more sheet residues than helix residues, making the summed PRS effectiveness larger for the sheet group. This analysis is a comparison within the set of known UBL family members analyzed here, not a statistical inference based on sampling from a larger population; results are therefore reported as relative rankings within the analyzed set.

**Figure 4.**
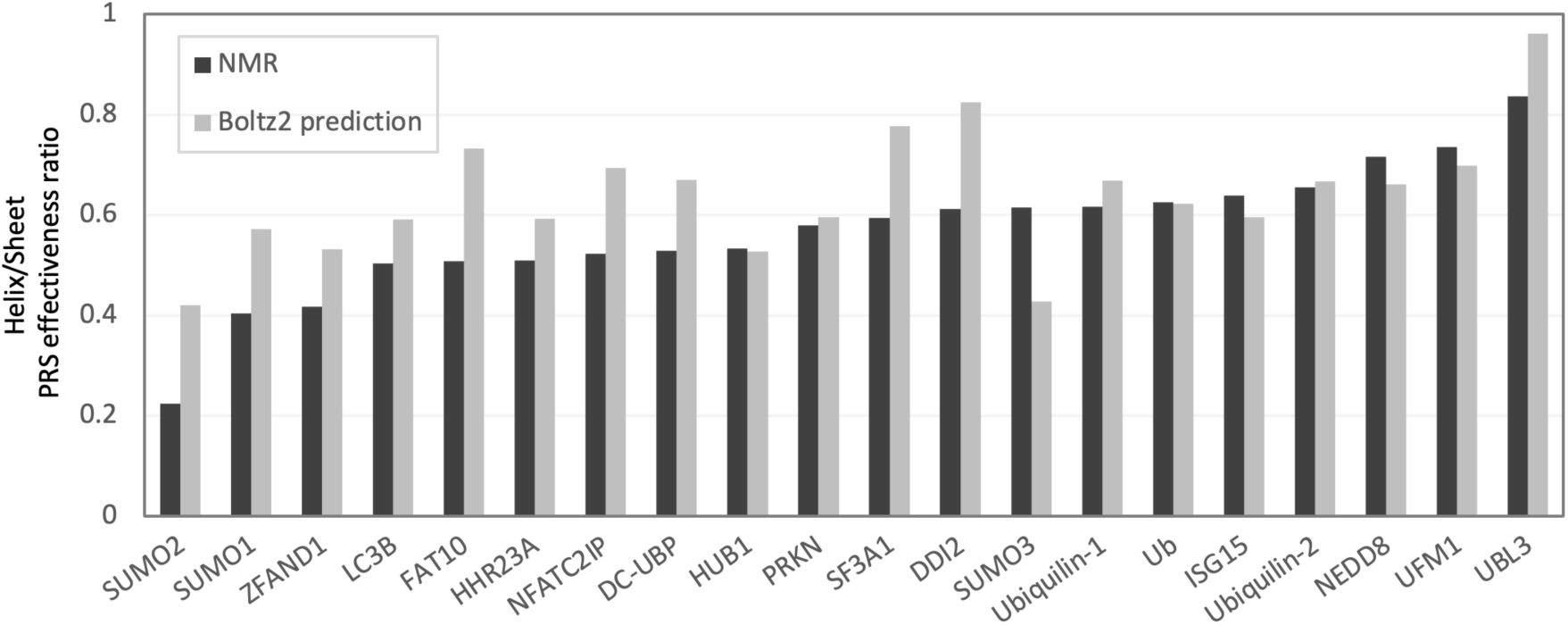
UBL3 exhibits the highest helix/sheet PRS effectiveness ratio among UBLs. Helix/sheet PRS effectiveness ratios of UBL domains derived from NMR structures and Boltz2-predicted models. Ratios represent the summed PRS effectiveness of helix residues divided by that of sheet residues within the UBL region. For NMR-derived data, the number of models listed in Table 1 was used; for Boltz2-predicted data, the mean across the 10 generated models was used.

To corroborate this finding with an independent approach, we performed the same analysis using Boltz2-predicted structures of the UBL domains of the same UBLs. Consistent with the NMR-based results, UBL3 again showed the highest helix/sheet PRS effectiveness ratio.

Although SUMO2/3 showed deviations between the NMR-derived and Boltz2-predicted datasets, the overall trend was preserved, with UBL3 exhibiting the highest helix/sheet PRS effectiveness ratio.

## CONCLUSION

This study provides the first systematic structural dynamics characterization of the UBL3 UBL domain. Using NMR ensemble analysis, ANM, and PRS, we demonstrate that C-terminal flexibility is a conserved yet variable property across UBL family members, while UBL3 uniquely exhibits the highest helix/sheet PRS effectiveness ratio among the 20 UBLs analyzed. These findings establish structural principles underlying UBL3’s broad interaction capacity, and identify helix residues as functionally important dynamic control sites that may represent rational targets for future pharmacological intervention.

## DISCUSSION

Analysis of the UBL3 NMR structure data consistently revealed pronounced structural diversity and high dynamic fluctuation in the C-terminal region. Proteins are now widely recognized to function not as single static structures but as dynamic ensembles of multiple structural states, and such structural dynamics are considered an essential element of allosteric regulation and functional expression [43]. Recent studies have shown that intrinsically disordered regions enhance the plasticity of interaction networks and increase binding diversity [44]. Protein internal flexibility and structural fluctuations are also closely linked to interaction modes and functional regulation; large-scale molecular dynamics database analyses have enabled systematic description of the distribution of flexibility and dynamic properties [45]. In particular, regions with low secondary-structure propensity and disordered regions contain sequence elements involved in binding and post-translational modification sites, enabling dynamic intermolecular interactions through conformational plasticity [46]. The C-terminal region of UBL3 contains the cysteine residue (C113) essential for post-translational modification and the CAAX motif (C114) involved in membrane anchoring. The dynamic nature of this region may confer the structural freedom required for interactions with modifying enzymes and membrane-anchoring factors.

Comparative analysis across UBLs further showed that high C-terminal fluctuation is not specific to UBL3 but is a dynamic feature shared by multiple UBLs, including ubiquitin (Ub) and NEDD8. This suggests that C-terminal flexibility within the UBL family is part of a plastic structural property tuned to the functional demands of each protein, rather than a conserved structural element.

PRS effectiveness analysis showed that perturbations applied to residues in the α-helix region of the β-grasp fold of UBL3 made high contributions to the collective motions of the entire protein. This is consistent with the concept that control sites within a network, rather than local flexibility per se, are the key determinants of protein functional regulation [47]. Recent allostery research also emphasizes the importance of collective structural behavior and dynamic coupling between domains in functional control [43]. PRS has additionally been applied to identify functionally important control residues and allosteric regulatory sites during conformational transitions, as exemplified by studies of structural rearrangements during caspase-6 activation [48]. A notable finding in the present study is that high-PRS-effectiveness residues in UBL3 were not strictly limited to direct binding interfaces or pocket residues. High-PRS-effectiveness residues are thought to act as network nodes that mediate structural changes and information propagation arising from allosteric perturbations [49]. Such residues may therefore function as control points governing conformational transitions and the distribution of motion, providing an important clue for understanding the relationship between dynamic structure and functional regulation.

Integrating these findings, the high C-terminal flexibility and helix-centered dynamic control within the β-grasp fold suggest a functional division of labor in UBL3: the C-terminus provides structural adaptability, while the central helix mediates long-range dynamic information transfer.

At the same time, elastic network model-based structural dynamics analyses, including PRS effectiveness analysis, efficiently extract the intrinsic collective motion properties of proteins based on low-frequency modes, but they rely on a coarse-grained model with a simplified force field and therefore do not fully reproduce the detailed atomic-level dynamic behavior or the experimentally observed fluctuations [50]. Given this premise, the fluctuating regions and high-PRS-effectiveness residues identified in the present study should be viewed as the structural potential of UBL3 in the free state.

Using both NMR-derived structural data and Boltz2-predicted structures as independent sources, UBL3 showed the highest helix/sheet PRS effectiveness ratio among the UBLs compared. This result indicates that the characteristic high dynamic contribution of helix structures in the UBL3 UBL domain is consistently maintained regardless of differences in structure determination method or prediction algorithm. In the present study, Boltz2-predicted structural models were primarily used as a validation dataset to verify the robustness of findings obtained from NMR-based structural dynamics analysis. Deep learning-based structure prediction methods such as Boltz2 and AlphaFold2 have recently been reported for use not only in highly accurate prediction of static three-dimensional structures, but also in diverse analyses including extraction of structural features, evaluation of mutational effects, and identification of functional sites [41]. These prediction methods primarily produce a single high-confidence structural model, however, and do not directly reflect the structural ensemble or fluctuations observed by NMR [51]; predicted structures thus provide a representative static picture but do not necessarily encompass the full landscape of dynamic diversity. The use of Boltz2 in this study was not intended for dynamic interpretation of the predicted structures themselves, but rather as an independent validation to confirm that the trends observed in NMR-based structural dynamics analysis are reproduced by an independent structure generation method.

Deep learning-predicted protein structures have been widely used not only as structural models per se but also as foundational information for functional analysis and molecular property evaluation. For example, expansion of the AlphaFold Protein Structure Database has made high-accuracy predicted structures available for vast numbers of protein sequences, enabling function annotation and analysis based on structural information [52]. Methods that use these predicted structures to identify binding sites and functional residues have also been developed, including binding site prediction based on local structural features [53]. Progress has also been made in interpreting the effects of disease-associated variants through protein stability evaluation and structural change analysis based on predicted structures, with application to variant functional assessment for BRCA1 [54]. A distinctive feature of the present study is the use of NMR-based structural dynamics analysis as the primary framework, with independent validation of the findings using deep learning-predicted structures. This complementary use of experimental and deep learning-predicted structures represents a useful approach for enhancing the reproducibility and generalizability of findings in structural dynamics research.

## DATA AVAILABILITY

All sequences and PDB entries used in this study are listed in the Materials and Methods section (Table 1).

## CODE AVAILABILITY

The code used for the analysis and figures in this study can be accessed at the following repository: https://github.com/TKahyo/PRS-ensemble-pipeline

## ACKNOWKEDGE

This work was supported by JSPS Grant-in-Aid for Scientific Research (C) (Grant numbers 24K09982), Grant-in-Aid for Scientific Research (B) (Grant Number JP22H02793) and Grant-in-Aid for Challenging Exploratory Research (Grant Number JP23K18197). This research is partially supported by the Recommendation Program for Young Researchers and Woman Researchers, Supercomputing Division, Information Technology Center, The University of Tokyo.

## DECLARATION OF INTERESTS

The authors declare that they have no known competing financial interests or personal relationships that could have appeared to influence the work reported in this paper.

## Notes

### Competing Interest Statement

The authors have declared no competing interest.

